# Volatile organic chemical analyser (eNose) diagnosis of dogs naturally infected with visceral leishmaniasis in Brazil

**DOI:** 10.1101/545202

**Authors:** ME Staniek, L Sedda, TD Gibson, CF de Souza, EM Costa, RJ Dillon, FGC Hamilton

## Abstract

Visceral leishmaniasis (VL) in Brazil is a neglected vector-borne tropical parasitic disease that is responsible for several thousand human deaths every year. The numbers of cases more than doubled between 1990 and 2016. Transmission occurs when sand flies become infected after feeding on infected dogs (the reservoir host) and then subsequently on humans. A major component of the VL control effort is the identification and euthanasia of infected dogs to remove them as a source of infection. Rapid, accurate identification of infected dogs would be key to this strategy.

Here we demonstrate the potential of a volatile organic chemical analyser (eNose) to rapidly and accurately identify dogs infected with *Leishmania infantum* parasites. The eNose could discriminate between the odours present in the hair of infected and uninfected dogs with greater than 95% sensitivity and 95% specificity. The device was sufficiently sensitive to be able to identify infected dogs even when parasite loads in the circulating blood were very low.

Future improvements to VOC analyser technology, portability and ease of use suggest that this methodology could significantly improve the diagnosis of VL infected dogs in Brazil and elsewhere and with other parasitic diseases such as Malaria, Chaga’s Disease and Leishmania in humans.

## Introduction

Visceral leishmaniasis (VL) is a neglected tropical disease caused by Protist parasites belonging to the genus *Leishmania*. Globally over 350 million people are at risk of infection with an estimated 200-400 thousand cases annually and an estimated 10% fatality rate. Ninety percent of all reported VL cases occur in only six countries including Brazil (Alvar et al, 2012; OPAS/WHO, 2017).

In Brazil, transmission of *Leishmania (Leishmania) infantum* (Kinetoplastida: Trypanosomatidae) occurs between domestic dogs *Canis familiaris* (Carnivora: Canidae) (the reservoir host) and from dogs to humans when an infected female sand fly vector *Lutzomyia longipalpis* (Diptera: Psychodidae) takes a blood meal.

Despite substantial efforts by the Brazilian Ministry of Health (MoH) the burden of VL in Brazil more than doubled between 1990 and 2016 (Martins-Melo et al., 2018). The increase is probably due to the spread of the vector into urban areas as a result of human migration into cities (Werneck, 2010) and the expansion of the range of the vector into new areas because of environmental degradation (Brazil et al., 2011; Salomón et al. 2015; Casanova et al., 2015). Given the spread of the disease and increase in cases it is also likely that current VL control measures are inadequate (Quinnell and Courtenay, 2009).

The control of VL in Brazil has three main components. Insecticides are applied in houses and animal sheds to lower the vector population density and reduce vector-human contact. Secondly, diagnosis and treatment of human cases to prevent severe forms of the disease and death. Finally, the identification and euthanasia of seropositive canine cases to decrease the sources of infection for the vector (MS/SVS/DVE 2014; Albuquerque e Silva et al, 2018).

Modelling predicts that the dog-culling program in Brazil should be effective in areas of low, medium but not high *Leishmania* transmission (Costa et al., 2013). However, the practice is controversial and despite the euthanasia of thousands of canines with suspected and confirmed infection each year the program has been unsuccessful (Dantas-Torres et al., 2012, Albuquerque e Silva et al, 2018). There are a number of possible explanations for this situation A). A shortage of qualified professionals caused by financial constraints leading to delays in collections, performance of routine diagnostic tests and subsequent removal of seropositive dogs. B). A failure to identify and remove the high proportion of asymptomatic animals C). Refusal of dog owners to comply with surveillance measures. D). A high rate of dog replacement with young immunologically naïve dogs. E). Lack of an accurate point-of-care diagnostic test (von Zuben and Donalísio 2016).

Identification of dogs infected with CVL follows a two stage serodiagnostic protocol recommended by the Brazilian MoH. Initial screening using the Dual-Path Platform (DPP^®^ CVL) immunochromatography diagnostic test is followed by a laboratory-based ELISA (EIE CVL) confirmatory test within 2 weeks. This protocol has an accuracy of 91% for symptomatic dogs and 97% for asymptomatic if applied in this way (Fraga et al, 2015). The DPP CVL test has been assessed several times since it was introduced. The most recent assessment showed that it has 75% sensitivity and 73% specificity for asymptomatic dogs and 94% sensitivity and 56% specificity for symptomatic dogs (Figueiredo et al, 2018) suggesting that the DPP test is better for confirming than identifying cases (Grimaldi et al, 2012).

The concept of volatile organic compounds (VOCs) as diagnostic aids to signal a disease is well established and since antiquity, many physicians have used odours associated with disease to help diagnose their patients (Liddell, 1976). Modern analytical techniques such as single ion flow tube mass spectrometry (SIFT-MS) and chemiresistive sensors have taken the concept to the point of widespread clinical application. Volatile markers from human breath can be used to identify a variety of disease states e.g. inflammatory bowel disease (IBD), chronic liver disease, diabetes, *Pseudomonas aeruginosa* infection and adenocarcinomas (Kumar et al., 2015; Smith and Španěl, 2016). A recent study has shown that the use of VOCs is sufficiently robust to discriminate between 14 cancerous and other disease states (Nakhleh et al, 2017).

Parasite infections of humans and animals also alter the odour of the host animal. The odours of golden hamsters infected with *Le. infantum* are more attractive to female sand flies than the odours of uninfected hamsters (O’Shea et al., 2002; Nevatte et al., 2017). The odour obtained from the hair of dogs infected with *Le. infantum* in Brazil was found to be significantly different to the odour of uninfected dogs. These odour differences which were detected by coupled gas chromatography-mass spectrometry (GC/MS) and multivariate statistical analysis indicated the increased presence of a small number of primarily low molecular weight aldehydes (octanal, nonanal), alkanes (undecane, heptadecane) and 2-ethylhexyl-salicylate (Oliveira et al., 2008; Magalhães-Junior et al., 2014). More recently, odours were also implicated in children infected with the infectious gametocyte stage of the malaria parasite *Plasmodium falciparum were* found to be more attractive to the mosquito vector *Anopheles gambiae* (Lacroix et al., 2005). This phenomenon occurred even when the gametocytemia was very low and was associated with changes in aldehyde concentration of the foot odours of the infected children (Busula et al, 2017; Robinson et al, 2018).

GC/MS analysis is a useful research tool but its use as a widely available diagnostic tool is unrealistic because of significant associated infrastructure and personnel costs. An alternative means of detecting the odour change associated with parasitaemia is required that would fulfil the majority of the World Health Organisation ASSURED criteria (affordable, sensitive, specific, user-friendly, rapid and robust, equipment free and deliverable to end-users) (Peeling et al, 2006) for all new point-of-care diagnostics tools. VOC analysers (eNoses) may fulfil WHO criteria, they can detect differences in the odours from sputum of tuberculosis (TB) infected and TB uninfected patients with sensitivity, specificity and accuracy of around 70% (Kolk et al, 2010). The aim of the present study was to determine if the odour of dogs naturally infected with *Le. infantum* could be detected using a commercially available VOC analyser.

## Materials and Methods

### Study Site

Governador Valadares (18°51′S, 41°56′W) (Minas Gerais State, Brazil), located in the valley of the Rio Doce 320 km northeast of Belo Horizonte is a city of approximately 280,000 people. The climate is temperate, characterised by dry winters and hot, wet summers (Alvares et al., 2013). Studies in Governador Valadares in 2013 found that an average of 30% of dogs from 16,529 samples taken from 35 urban and rural districts were seropositive for canine visceral leishmaniasis (CVL) (Barata et al., 2013). From 2008 until 2017, 193 human VL cases were recorded in Governador Valadares with a fatality rate of 14.5%. (Unpublished data provided by the Governador Valadares Department of Epidemiology).

### Ethics

Dog blood and hair samples were taken from dogs and were microchipped with the informed consent of their owners. Ethical approval was obtained from the Comissão de Ética no Uso de Animais (CEUA), Instituto Oswaldo Cruz (licence L-027/2017) in Brazil and Lancaster University Animal Welfare and Ethics Review Board (AWERB) in the UK. The CEUA approval complies with the provisions of Brazilian Law 11794/08, which provides for the scientific use of animals, including the principles of Brazilian Society of Science in Laboratory Animals (SBCAL). The AWERB approval complies with the UK Home Office guidelines of the Animals in Science Regulation Unit (ASRU) and in compliance with the Animals (Scientific Procedures) Act (ASPA) 1986 (amended 2012) regulations and was consistent with UK Animal Welfare Act 2006.

### Dog recruitment

A 2 year cross-sectional study in the Altinopolos district of Governador Valadares was initiated in August 2017 by initial recruitment and sampling of 185 dogs. The area was chosen because of the high prevalence of CVL (average incidence 33.8%) (Barata et al., 2013) and the large population of household-owned dogs (ca. 2000) (Centro de Controle de Zoonoses (CCZ) survey) located there. The dogs were microchipped to aid their identification. Inclusion criteria: dogs aged ≥ 3 months, dogs without previous clinical assessment or laboratorial diagnosis for CVL. Exclusion criteria: pregnant/lactating bitches; aggressive dogs; stray dogs. In April 2018 149 dogs were sampled, this number included 133 dogs that were resampled from the 2017 cohort and an additional 16 from the CCZ facility that had been assessed as infected to increase the proportion of samples from infected dogs.

Between 5ml and 10ml of peripheral blood was collected in 10ml K2 EDTA-coated tubes (BD Vacutainer, UK) via cephalic or jugular venepuncture by a qualified vet in 2017 and by a CCZ qualified phlebotomist in 2018. Samples were placed in containers marked with the microchip bar code to aid subsequent tracking and identification. Blood samples were stored in a cool box with a freezer pack before being transferred to a fridge (4°C) prior to processing.

Hair samples were obtained by cutting the dorsal hair close to the skin using surgical scissors that had been washed with hexane prior to the collection of each sample by members of the LU research team. A minimum of 2g of hair was collected from each dog. All hair samples were placed in individual foil bags (110mm x 185mm; Polypouch UK Ltd, Watford, England) heat sealed and stored at 4°C prior to analysis.

All dogs were assessed for clinical signs of *Leishmania* infection by veterinarians and CCZ CVL control specialists. The animals were classified according to the presence of clinical signs which were recorded for each dog. The main signs of CVL considered were onychogryphosis, ophthalmologic abnormalities, adenitis, cachexia, hepatosplenomegaly, alopecia, crusted ulcers and lesions; dogs were classified as asymptomatic (the absence of clinical signs), oligosymptomatic (the presence of one to three clinical signs), or symptomatic (the presence of more than three clinical signs (Mancianti et al., 1988).

### Molecular Diagnosis of Dogs

#### DNA Extraction

Collected blood samples were centrifuged at 2500 x g for 10 minutes at room temperature and the top layer of buffy coat removed, placed in 1.5ml Eppendorf tubes and stored at −20°C until DNA extraction. The DNA was extracted from 200μl of buffy coat samples using the QIAamp DNA Blood Mini Kit following the manufacturer’s instructions. Cell lysis was mediated using protein kinase with a final elution volume of 50μl.

#### Qualitative detection of Leishmania DNA

Conventional PCR was initially used to ascertain which blood samples were positive for *Leishmania infantum*.

Following primer optimization, extracted DNA from canine blood obtained during August 2017 and April 2018 were tested using Primer pair MaryF (5’ – CTT TTC TGG TCC TCC GGG TAG G – 3’), and MaryR (5’-CCA CCC GGC CCT ATT TTA CAC CAA – 3’ (Mary et al, 2004). The reactions were performed in a final volume of 25µl containing 0.5µl DNA template (100ng µl-1), 12.5µl Mastermix (dH_2_O, Buffer 5x, MyTaq redmix polymerase, dNTP’s) and 10µM of each primer. The PCR amplifications were performed in a TECHNE^®^ Prime Thermal Cycler (Cole-Palmer Ltd., Staffordshire, UK) using the following conditions: 95°C for 5mins and 30 cycles of 95°C for 30sec, 57°C for 30sec and 72°C for 60sec, followed by 72°C for 10min.

The PCR products were analysed by gel electrophoresis using 2% agarose gels run at 90V for 1hr 30 minutes and visualized under UV light following the addition of 6.5µl of 10,000x SYBR Safe (Thermo Fisher Scientific, UK) to each gel. Samples were run 3 times and dogs were considered to be infected if 2 or 3 out of the 3 replicates were positive.

#### Quantitative detection of Leishmania DNA

A real-time quantitative PCR (qPCR) for detection and quantification of *Le. infantum* DNA in positive dog samples from both sampling occasions (August 2017 and April 2018) was performed using MaryF/R primers.

The qPCR amplifications were performed on a Bio-Rad C1000™ Thermal Cycler with each reaction consisting of a final volume of 13.0µl; 12.0µL of PCR mix plus 1µL of DNA (approximately 75-100 ng/µl per reaction). The qPCR mix was composed of 6.25 µL 2x QuantiNova SYBR Green PCR Master Mix, 0.5 µL of each primer (MaryF/R, corresponding to 10 mmol) and 4.75 µL of water (Costa Lima Junior *et al* 2013).

The amplification was performed in triplicate (Ceccarelli *et al*, 2014) at 94°C for 10 min, followed by 40 cycles at 94°C for 30 sec, 60°C for 20 sec and 72°C for 20 sec. At the end of each run, a melt curve analysis was performed from 55°C to 95°C in order to identify the formation of non-specific products as well as primer dimers. A standard curve was established using extracted *Le. infantum* DNA; 1:10 serial dilutions, ranging from 10,000 to 0.01 parasites per ml.

#### VOC Analysis

Initial VOC analysis was carried out on all (n=11) of the infected dog hair samples and a sub-set of the uninfected dog hair samples (n=44) collected in 2017. The uninfected dogs were selected from groups of dogs matched by shared characteristics (age, sex and whether or not treatment for ectoparasites was received) with infected dogs (Table 1). Subsequently the VOC analysis was carried out on all the infected dog hair samples (n=44, including 10 CCZ infected dogs) and all of the uninfected dog hair samples (n=105, including 6 CCZ uninfected dogs) collected in 2018.

**Table 1.** *Leishmania infantum* infection status of dogs sampled in Governador Valadares, Minas Gerais.

| Month of sample collection | Source of dogs | Number of dogs sampled | Leishmania positive dogs (PCR) |  |  | Prevalence (%) |
| --- | --- | --- | --- | --- | --- | --- |
|  |  |  | symptomatic | oligosymptomatic | asymptomatic |  |
| Aug-17 | Altinópolis | 185 | 0 | 3 | 8 | 6.0 |
| Apr-18 | Altinópolis | 133 | 3 | 7 | 24 | 25.6 |
| Apr-18 | CCZ GV | 16 | 6 | 2 | 2 | 62.5 |

Dogs from Altinópolis, Governador Valadares sampled in August 2017 (n=185) and April 2018 (n=133). An additional 16 dogs, considered positive by CCZ, were sampled in April 2018. PCR experiments were performed in triplicate with both positive and negative dogs identified.

A VOC analyser (Model 307, RoboScientific Ltd, Leeds, UK) with 11 functioning semi-conducting polymer sensors was used for the analysis. Each sensor has 2 outputs (positive and negative) giving a total of 22 responses. Two calibration points were automatically set by the sensor unit; the first was the baseline obtained when carbon-filtered air was passed over the sensor at a flow rate of 200ml min^-1^ which was automatically adjusted to zero on the Y-axis scale, and the second was a reference point obtained from sampling the head space of 5ml of a liquid water control in a plastic vial.

The chemical sensors were thin films of semi-conducting polymers deposited onto interdigitated gold structures on a silicon substrate. We used 12 different sensor types chosen from a group of polymers that included polyaniline, polythiophene and polypyrrole. Each sensor had semi-selectivity to a different group of volatile chemicals; aldehydes, alcohols, amines, organic acids and ketones etc. In this way a digital fingerprint of the VOC mixtures emanating from the samples was generated. Two similar sensor arrays were used in the study, the second array being a derivation of the first with 50% of the sensors being identical to the first array.

The interaction of the mixtures of VOCs in the samples with the semi-conducting polymer surfaces produced a change in electrical properties (e.g. voltage and resistance) over time. This change was measured, recorded and simultaneously displayed on the VOC analyser data logger screen for each sensor. Four parameters were used from each sensor response; the divergence from the baseline (maximum response), the integrated area under each response curve, absorbance and desorbance. Therefore, the total number of VOC measurements produced for each sample were 88 (11 sensors x 4 parameters and 2 outputs – positive or negative). The sampling profile was set at 2 seconds baseline, 7 seconds of absorption, a 1 second pause, 5 seconds desorption and 12 seconds flush to bring the sensors back to baseline.

Water (DD;10µl) was injected into each foil bag containing the dog hair samples with a Hamilton syringe and inflated with 140ml of laboratory air using a diaphragm pump. The samples were then incubated at 50°C for 15 minutes in an oven, then allowed to cool to room temperature for 5 minutes prior to head space analysis.

For the analysis each foil bag was sampled by insertion of an 18-gauge needle connected to a PTFE tube through the sidewall of the bag with the tip placed into the head space of each bag. This was connected to the sample port of the VOC analyser and the head space sample was therefore passed over the 12 sensor surfaces. The original flow rate for the sampling was 200 ml min^-1^. The headspace of each foil bag was sampled 4 times. The first sample was disregarded as potentially it could contain volatile carryover from the previous sample and thus, we retained the data from the next 3 samples for analysis. The individual dog hair samples in each experiment were tested randomly with each sample used once only.

### Data Analysis

To test the ability of the VOC analyser to differentiate between the odours of infected and uninfected dogs, we employed a statistical program, MCLUST (Fraley and Raftery 1999), a model-based clustering and classification algorithm embedded in R-cran statistical software (R-cran, R Core Team 2018). This was applied to the known data classes (infected or uninfected dogs). The initial analysis indicated that the data was overfitted, therefore we divided the infected and uninfected dog classes into sub-classes and the analysis was repeated. The robustness of the classification was then evaluated by two types of cross-validation; out-of-sample cross validation (CV) and confounder cross validation (CCV). Finally, the importance of each variable produced by the VOC analyser in discriminating between the infected and uninfected sub-classes was assessed by variable permutation analysis. A more detailed explanation of the rationale for this analysis approach is provided in the Supplementary Material 1 (Analysis Rationale) and a more extensive description of the algorithms is provided in (Scrucca, Fop et al. 2016).

The VOC analysis dataset contained data from:

a) Infected and uninfected dog odour collected in 2017.
b) Infected and uninfected dog odour collected in 2018 including samples collected from CCZ dogs.

Three replicate VOC analyser readings were obtained for each dog odour sample. These replicates were considered to be independent, i.e. the three VOC replicates for each dog were considered as coming from three different dogs (a common procedure in clustering).

The analysis aimed to identify any significant differences in the VOC analyser variables (used to obtain the means and covariances of the infected and uninfected classes and/or sub-classes) of infected and uninfected dogs so as to be able to accurately predict the infection state of newly sampled dogs. Initially, the data was evaluated to determine 1. if infected and uninfected dogs in both 2017 and 2018 could be statistically separated and 2. if the uninfected dogs in 2017 were statistically separate from uninfected dogs in 2018.

To take account of overfitting (Bettenburg, Nagappan et al. 2015), we reclassified the infected and uninfected classes into sub-classes using the function Mclust. The optimal inferential method and number of subclasses for infected and uninfected classes was obtained by Bayesian information criterion (BIC) (Supplementary Material 1 (Analysis Rationale)), bootstrapping and the likelihood ratio test (function MClustBootstrapLRT).

### Cross-validation (CV) and confounder cross-validation (CCV) analysis

Once the best model (number of subclasses and model components) had been found, we tested for the importance of the variables in clustering by permutation analysis; while the predictive capacity of the model by using leave-one-out cross validation (CV); and finally, the capacity of the model to recognise sample confounders by developing a technique named confounder cross-validation (CCV). For the latter 10% of the data from the infected class were placed in the uninfected class and vice versa (leaving the remaining 90% in their correct class for training in both cases). These analyses were done by compiling algorithms that included some of the MCLUST components (Mclust, MclustDA, predict) and permutation functions.

### Results: Molecular diagnosis

#### Qualitative detection of Leishmania DNA

PCR revealed that 11/185 (6%) dogs were positive for *Le. infantum* infection in August 2017 and 34/133 (26%) in April 2018 (Table 1) representing a 20 percent increase in infection rate over the 8-month period between sampling points. The typical PCR results showed a band at 140bp of varying intensities representing a semi-quantitative indication of parasite presence in individual samples (Figure 1).

**Figure 1.**
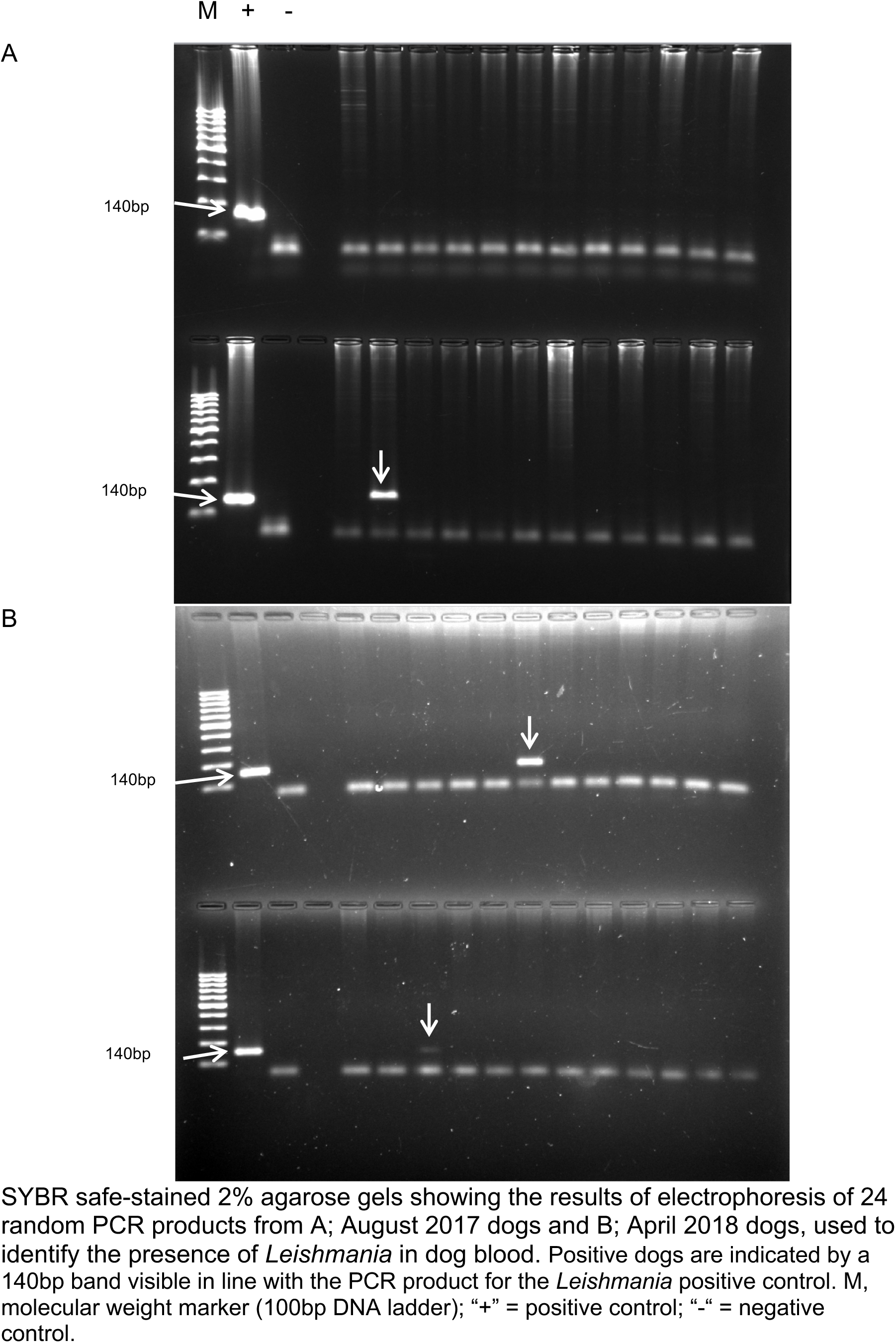
Preliminary detection of *Leishmania infantum* in dog blood samples

In 2017, 3 out of the 11 positive dogs presented as oligosymptomatic and 8 were asymptomatic. In 2018, 3 dogs were symptomatic, 7 were oligosymptomatic and 24 were asymptomatic (Table 1). Of the 174 uninfected 2017 dogs 42 were lost to follow-up in 2018. There were 7 oligosymptomatic dogs and 3 had become symptomatic in the remaining 133 dogs. In the 2017 cohort 55% of the infected dogs had 3 out of 3 positive PCR results and 45% had 2 out of 3 positive PCR results. In the 2018 field collected cohort 47% had 3/3 + PCR results and 53% had 2/3 + PCR results. In the 2018 CCZ collected cohort 80% had 3/3 + PCR results and 20% had 2/3 + PCR results.

The evaluation indicated that the most frequently occurring clinical signs were skin lesions including dermatitis (18% 2017; 27% 2018) and ulcerative lesions (0% 2017; 25% 2018), long nails (9% 2017; 25% 2018) and signs of conjunctivitis (18% 2017; 14% 2018).

PCR diagnosis of the 16 CCZ dogs, sampled in April 2018, that were assumed to be VL infected, indicated that 10 (63%) were positive and the remaining 6 cases were not infected.

### Quantitative detection of Leishmania DNA

The kDNA qPCR assay showed that parasite loads ranged from 0.4 to 103 parasites ml^-1^ in 2017 and from 1 to 850 parasite ml^-1^ in 2018 (both field and CCZ collected) (Figure 2).

**Figure 2.**
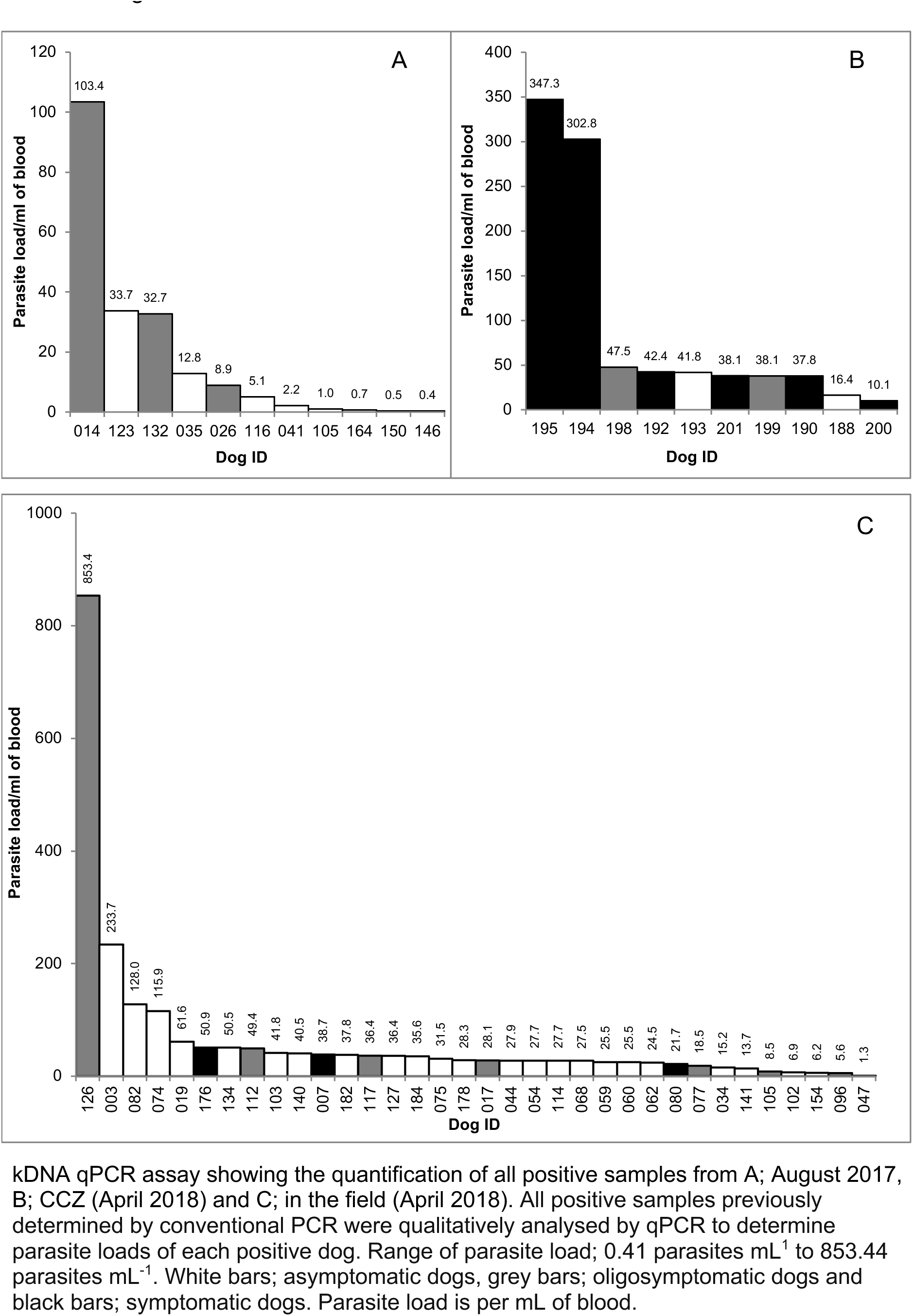
Quantitative estimation of *Leishmania infantum* in blood samples from infected dogs.

This large variation in parasitic load over the study period can be observed through analysis of the median values which ranged from 5.06 parasites/ml (dog #116) in 2017 to 28.32 parasites/ml (dog #178) in 2018. Comparisons of parasitic load among the samples revealed that dog #126 in 2018 (853.44 parasites/ml) exhibited the highest degree of parasitism with dog #146 in 2017 (0.41 parasites/ml) exhibiting the lowest.

The average value of CT (ΔCT) obtained for dog #126 in 2018 (highest degree of parasitism) and for dog #146 in 2017 (lowest degree of parasitism) were the following: dog #126; 20.43 and dog #146; 30.45. A lower CT value correlates with a higher parasitic load per ml of blood.

### Data Analysis

The best model for the analysis of all infected and uninfected dog classes was EEE apart from the uninfected 2017 dogs which was VVI (Scrucca, Fop et al. 2016). The EEE model assumes ellipsoidal covariances and equal shape, volume and orientation for all the classes. The VVI model assumes diagonal covariances with orientation parallel to the coordinate axes with variable shape and volume for the classes (Fraley and Raftery 2007). Between 14 x 9 classes were assessed (i.e. 126 mixture models) (Andrews and McNicholas 2012) (Supplementary Information Table S1 (2017 data) and S2 (2018 data)). Clustering analysis of 2017 dogs identified 1 class for uninfected dogs and 3 classes for infected dogs and for the 2018 dogs 2 classes for uninfected dogs and 6 classes for infected dogs were identified.

Confusion matrices of the separation obtained from the training set of uninfected vs infected dogs in 2017 and uninfected vs infected dogs in 2018 without sub-classes are given in Table 2A and 2C respectively and with sub-classes in Table 2B and 2D respectively below.

**Table 2.**
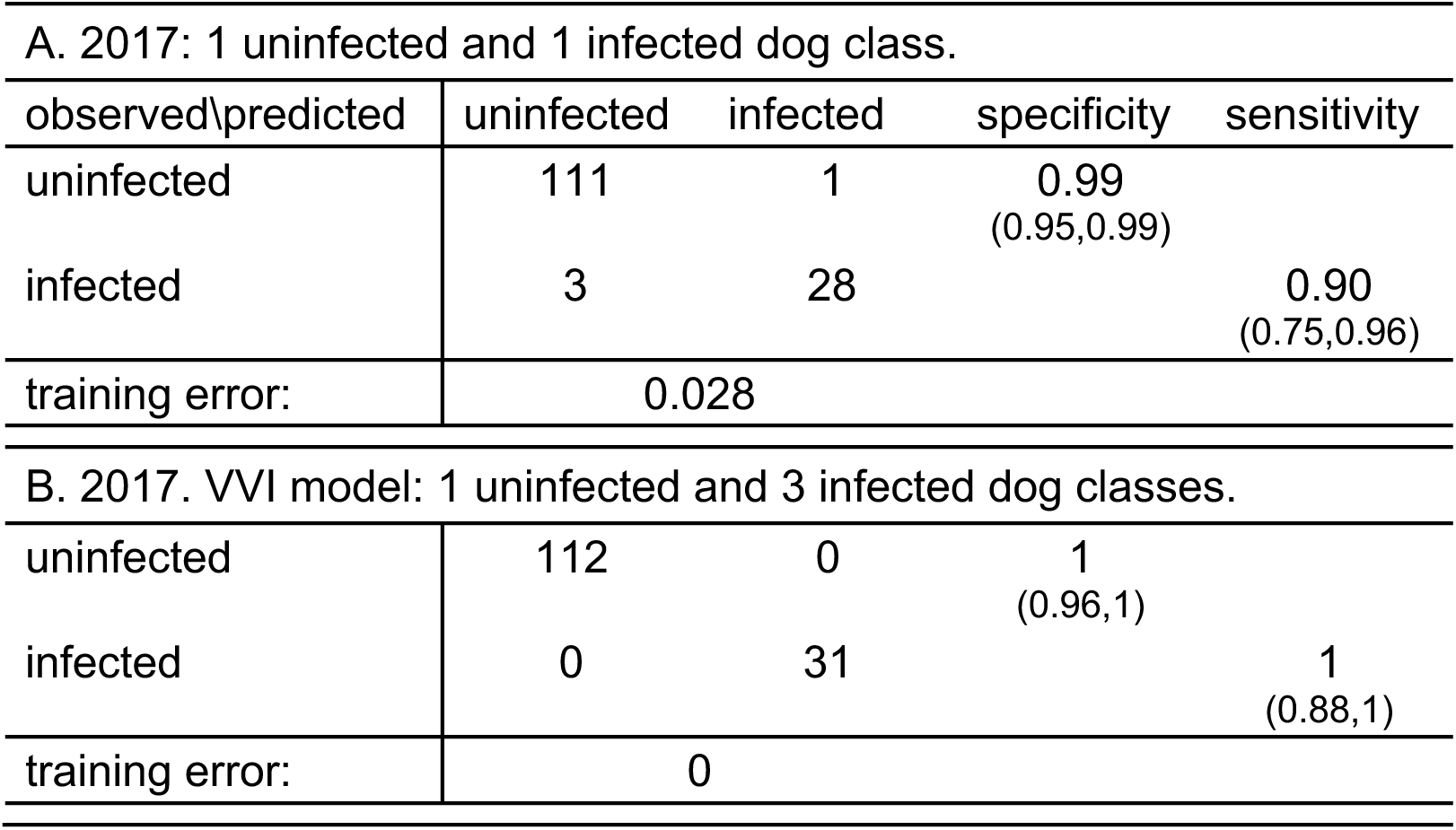
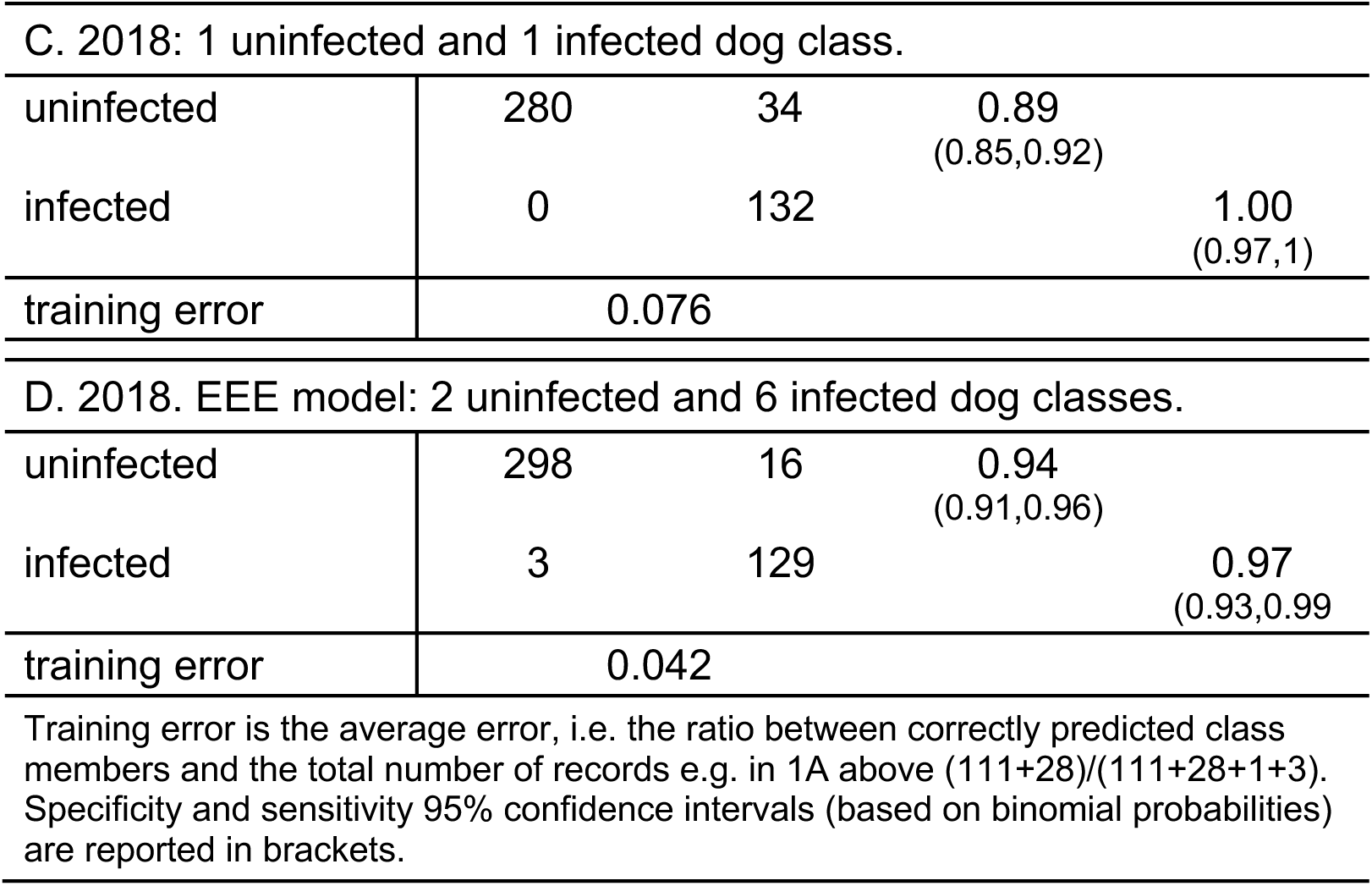
Confusion matrices for Gaussian mixture model EDDA classification.

These data show that in both years the infected dog odours were significantly different from the uninfected dog odours. In 2017 uninfected dogs were discriminated with 96% specificity and 97% sensitivity, that was improved to 100% for both metrics when the data was divided in sub-classes. The overall training error was reduced from 2.8% to 0%) when the 2017 data was divided in sub-classes.

In 2018 uninfected and infected dogs were discriminated with 89% specificity and 100% sensitivity and that was improved to 94% specificity and 97% sensitivity when the data was divided in sub-classes. The overall training error was reduced (from 7.6% to 4.2%) when the 2018 data was divided in sub-classes.

### Cross-validation (CV) and confounder cross-validation (CCV) analysis

The CV analysis returned a reduced sensitivity of 50% and specificity of 84% for 2017 dogs, and 48% sensitivity and 96% specificity for 2018 dogs (Table 3A and 3B first line and first two columns) indicating a reduced capacity to estimate true positives compared to the training set (as reported in Table 2) clearly showing model overfitting when considering 1 class for infected and 1 class for uninfected dogs. This was also confirmed by the CCV analysis, which was unable to find false positive and false negatives in the training groups. However, when the analyses were repeated on the EDDA models with sub-classes, both cross validation (CV) and confounder cross validation (CCV) calculations of sensitivity and specificity improved substantially (Table 3A and 3B second line). In other words, by identifying sub-classes for infected and uninfected dogs it was possible to obtain a better delineation of the multivariate space with improved predictivity capacity (CV analysis) and recognition of false positive and false negative in the training sets (CCV analysis).

**Table 3.** Comparison of sensitivity and specificity after CV and CCV analysis.

| A | 2017 dog samples |  |  |  |
| --- | --- | --- | --- | --- |
| Model | CV sensitivity | CV specificity | CCV sensitivity | CCV specificity |
| 2 classes | 0.50<br>(0.40,0.59) | 0.84<br>(0.75,0.89) | 0.11<br>(0.09,0.13) | 0.60<br>(0.56,0.62) |
| 4 classes | 0.75<br>(0.65,0.82) | 0.80<br>(0.71,0.86) | 0.60<br>(0.56,0.62) | 0.70<br>(0.67,0.72) |
| 2 classes = 1 uninfected + 1 infected class |  |  | 4 classes = 1 uninfected + 3 infected classes |  |
| B | 2018 dog samples |  |  |  |
| 2 classes | 0.48<br>(0.38,0.57) | 0.96<br>(0.90,0.98) | 0.18<br>(0.15,0.20) | 0.67<br>(0.64,0.69) |
| 8 classes | 0.93<br>(0.86,0.96) | 0.92<br>(0.85,0.95) | 0.74<br>(0.71,0.76) | 0.84<br>(0.81,0.86) |
| 2 classes = 1 uninfected + 1 infected |  |  | 8 classes = 2 uninfected + 6 infected |  |
| Specificity and sensitivity 95% confidence intervals (based on binomial probabilities) are reported in brackets. CV- cross validation; CCV- confounder cross validation. |  |  |  |  |

The e-nose variables important in the clustering are shown in Supplementary Information Table S3 (2017 data) and S4 (2018 data). A 0.99 P-value indicates that in 99% of the permutations the number of optimal clusters changed, indicating a strong influence of the variable in the final clustering.

## Discussion

The results presented in this study show that a VOC analyser can identify dogs infected with *Le. infantum* by analysis of their odour with very high sensitivity and specificity, regardless of parasite load or the presentation of clinical symptoms. We observed this outcome in two data sets from dog hair samples collected in 2017 (99% specificity and 90% sensitivity) and in 2018 (89% specificity and 100% sensitivity). When the small size of both data sets (2017, 55 dog hair samples: 2018, 149 dog hair samples) and consequent potential for overfitting was accounted for by improving the mixture of the models, both sensitivity and specificity increased (2017; 100% specificity, 100% sensitivity: 2018; 94% specificity, 97% sensitivity).

The robustness of the models was further tested by cross-validation and confounder cross validation analyses. The model predicted poorly when using 2 classes (infected and uninfected). However, when we accounted for the heterogeneity within each of these classes and subdivided them into either 4 subclasses (2017 data) or 8 subclasses (2018 data) the sensitivity and specificity and their confidence intervals improved significantly. In both cases the models accurately placed “unknown” dog odours in the correct infected or uninfected class with a high degree of specificity and sensitivity (Table 3 i.e. the first rows in the first 2 columns compared to the second rows first 2 columns).

The results suggested that the VOC analyser response was not related to the parasite load in the dog peripheral blood. As the analysis gave sensitivity and specificity responses substantially better than 90%, the effect of parasite load on the VOC analyser response is unlikely to have been significant as the majority of infected animals, regardless of parasite load were detected. However, determining the limits of detection will be important in the future.

Previous work has suggested that symptomatic dogs with a greater parasite load produced greater quantities of volatiles than infected asymptomatic dogs (Magalhães-Junior et al., 2014). However, asymptomatic dogs can contribute to disease transmission and VL control strategies should target infectious dogs rather than infected dogs *per se* and in particular the super-spreaders in the population (Courtenay et al., 2013; Nevatte et al., 2016). In this study we identified *Leishmania* DNA in circulating blood obtained by cephalic and jugular venepuncture, however the relationship between numbers of circulating parasites in peripheral blood and the infection status of the dog is unclear. In future studies, the skin parasite load, which appears to be more closely related to infectiousness (Courtenay et al, 2013) could be correlated with the odour profile.

It has been suggested that in dogs infected with *Le. infantum* the change in odour might be related to the immune response Magalhães-Junior et al., 2014. The results presented here suggest that the change in odour profile started to occur soon after infection as infected dogs with very low parasite loads could be differentiated from uninfected dogs. Changes in odour profile have been observed in other disease states where changes in relatively low molecular weight compounds were expressed as distinct and immediate changes arising from pathophysiological processes occurring and altering the body’s metabolism (Nakhleh et al., 2017).

Our results also suggest that there was no relationship between clinical state of infection (symptomatic, oligosymptomatic and asymptomatic) and detector response. The analyser could accurately detect asymptomatic dogs with low parasite levels as well as symptomatic dogs with high parasite loads.

It has been proposed that manipulation of the hosts chemical communication system could enhance the transmission of the parasite to the insect vector and potentially have a significant effect on the epidemiology of the disease (Hamilton and Hurd, 2002; O’Shea et al, 2002; Cator, 2017). Our study looked at volatile odours present on the dog hair only, it did not consider the effect of other volatiles, semi-volatiles and non-volatiles from other sources e.g. breath compounds, specialized scent gland secretions, sweat, urine or feces (Drea et al, 2013). The source of the odours that were detected by the VOC analyser is not clear, they could have arisen from the skin, as a result of the metabolic activity of skin microbiota (Ezenwa and Williams, 2014), the immune response or potentially directly from the *Le. infantum* parasites.

Our study did not examine the effect of other infections and the ability of the VOC analyser to differentiate between dogs infected with *Le. infantum* and other *Leishmania spp.* or other infections was not determined. In Governador Valadares dogs infected with *Le. amazonensis* have been found (Valdiva et al, 2016) and the sand fly vector *Lu. longipalpis* infected with multiple *Leishmania spp.* have been also been found (Cardoso et al, 2018) indicating that the epidemiological features require further work.

The application of VOC analyser technology is potentially a significant step towards the application of volatile odour analysis in diagnosis of parasitic disease. It raises the possibility that in the future a modified VOC device could provide a rapid, accurate, non-invasive point-of-care diagnostic tool for the specific diagnosis of leishmaniasis in dogs and humans. In our study we found that a small proportion of the sensor variables (2 out of 88 in 2017 and 3 out of 88 in 2018) contributed significantly to the outcome. Therefore, there is considerable scope for enhancing the sensitivity and specificity of the device through modifications to the sensor chemistry as well as incorporating further improvements to the field collection and analysis of odour. As well as further developments in robustness, portability and simplicity of the device all of which would improve the reliability and utility of the device in the field.

A reliable, rapid, accurate, non-invasive additional POC test that identifies *Leishmania* infection using a different set of disease markers in addition to the DPP CVL test could potentially eliminate the need for the relatively expensive, laborious in-laboratory ELISA confirmatory test that currently fails to rapidly diagnose and remove infected dogs from the population. The ability of a VOC analyser, that conforms to the WHO ASSURED criteria, to recognise infection in asymptomatic dogs and dogs with low levels of infection would be of great benefit to VL control programs. Further work to compare the sensitivity and specificity of a VOC test combined with DPP CVL diagnostics against DPP CVL combined with ELISA is required. The development of a non-invasive POC diagnostic tool based on host odour opens up a myriad range of opportunities to diagnose *Leishmania* infections in humans and other diseases such as malaria, trypanosomiasis and Chaga’s disease.

## Acknowledgements

We are grateful to CCZ in GV for their permission and practical support to help carry out the work. Ricardo Alves Miranda phlebotomist (CCZ, Governador Valadares, MG) for collecting the dog blood samples in 2018. Ms M. Bell Lancaster University for providing technical support. Prof P. Bates (Division of Biomedical and Life Sciences, Lancaster University) who supplied *Leishmania infantum* promastigotes (strain M4192). We are profoundly grateful to the dog owners of Altinopolos, GV for allowing us to carry out this study on their cherished companion animals.

## Author contributions

M.E.S., T.D.G., E.M.C., C.F.dS., and R.J.D. acquired the data; L.S., M.E.S., and T.D.G. analysed the data; L.S., J.G.C.H., M.E.S. and R.J.D. interpreted the data; J.G.C.H., L.S., M.E.S. and R.J.D. drafted the paper. J.G.C.H. conceived and designed the study and obtained research funding. All authors edited and approved the final version of the manuscript.

## Corresponding author

Correspondence to James G.C. Hamilton

## References

Albuquerque e Silva, R, de Andrade, AJ, Quint, BB, Raffoul, GES, Werneck, GL, Rangel, EF, Romero, GAS. (2018) Effectiveness of dog collars impregnated with 4% deltamethrin in controlling visceral leishmaniasis in Lutzomyia longipalpis (Diptera: Psychodidade: Phlebotominae) populations. Mem Inst Oswaldo Cruz, Rio de Janeiro, Vol. 113(5): e170377.

Alvar J, Velez ID, Bern C, Herrero M, Desjeux P, Cano J, et al. (2012) Leishmaniasis worldwide and global estimates of its incidence. PLoS ONE. 7(5):e35671. Epub 2012/06/14. pmid:22693548

Alvares CA, Stape JL, Sentelhas PC, de Moraes G, Leonardo J, Sparovek G. Köppen’s climate classification map for Brazil. Meteorologische Zeitschrift. 2013;22(6):711–28.

Andrews, J. L. and P. D. McNicholas (2012). “Model-based clustering, classification, and discriminant analysis via mixtures of multivariate t-distributions.” Statistics and Computing 22(5): 1021–1029.

Barata, R.A., Peixoto, J.C., Tanure, A., Gomes, M.E., Apolinário, E.C., Bodevan, E.C., de Araújo, H.S., Dias, E.S., Pinheiro Ada, C. (2013). Epidemiology of visceral leishmaniasis in a re-emerging focus of intense transmission in Minas Gerais State, Brazil. Biomed Res Int. 2013: 405083. https://doi.org/10.1155/2013/405083

Bell MJ, Sedda L, Gonzalez MA, de Souza CF, Dilger E, Brazil RP, Courtenay O Hamilton JGC. (2018) Attraction of Lutzomyia longipalpis to synthetic sex-aggregation pheromone: effect of release rate and proximity of adjacent pheromone sources. PLoS Negl Trop Dis 12(12): e0007007. https://doi.org/10.1371/journal.pntd.0007007

Bettenburg, N., M. Nagappan and A. E. Hassan (2015). “Towards improving statistical modeling of software engineering data: think locally, act globally!" Empirical Software Engineering 20(2): 294–335.

Brazil, RP, Brazil-Rocha, U, Brazil, BG (2011) Impact of climatic changes and habitat degradation on Phlebotominae (Diptera: Psychodidae) distribution and leishmaniases dispersion in Brazil. 7th International Symposium on Phlebotomine Sand flies 25–30 April 2011, Turkey.

Busula AO, Bousema T, Mweresa CK, Masiga D, Logan JG, Sauerwein RW, Verhulst NO, Takken W, de Boer JG. (2017) Gametocytemia and attractiveness of Plasmodium falciparum Infected Kenyan children to Anopheles gambiae mosquitoes. The Journal of Infectious Diseases, 216(3),291–295, https://doi.org/10.1093/infdis/jix214

Cardoso MS, Bento GA, de Almeida LV, de Castro JC, Cunha JLR, de Araújo Barbosa V, de Souza CF, Brazil RP, Valdivia HO, Bartholomeu DC (2018) Detection of multiple circulating Leishmania species in Lutzomyia longipalpis in the city of Governador Valadares, southeastern Brazil waiting full ref

Casanova C, Colla-Jacques FE, Hamilton JGC, Brazil RP, Shaw JJ. (2015) Distribution of Lutzomyia longipalpis Chemotype Populations in São Paulo State, Brazil. PLoS Negl Trop Dis. 9(3):e0003620. doi: 10.1371/journal.pntd.0003620.

Cator L. (2017) Editorial: Host Attractiveness and Malaria Transmission to Mosquitoes. The Journal of Infectious Diseases, 216(3), 289–290. https://doi.org/10.1093/infdis/jix216

Ceccarelli M, Galluzzi L, Migliazzo A, Magnani M. (2014) Detection and Characterization of Leishmania (Leishmania) and Leishmania (Viannia) by SYBR Green-Based Real-Time PCR and High Resolution Melt Analysis Targeting Kinetoplast Minicircle DNA. PLoS ONE 9(2), e88845. doi:10.1371/journal.pone.0088845

Costa DNCC, Codeço CR, Silva MA, Werneck GL. (2013) Culling dogs in scenarios of imperfect control: realistic impact on the prevalence of canine visceral leishmaniasis. PLoS Negl Trop Dis. 7(8): e2355.

Costa Lima Junior MS, Zorzenon DCR, Dorval MEC, Pontes ERJC. (2013) Sensitivity of PCR and real-time PCR for the diagnosis of human visceral leishmaniasis using peripheral blood. Asian Pac J Trop Dis. 3(1), 10–15.

Courtenay O, Carson C, Calvo-Bado L, Garcez LM, Quinnell RJ (2013) Heterogeneities in Leishmania infantum Infection: Using Skin Parasite Burdens to Identify Highly Infectious Dogs. PLoS Negl Trop Dis 8(1): e2583. doi:10.1371/journal.pntd.0002583

CCZ (2016) http://www.prefeitura.sp.gov.br/cidade/secretarias/saude/vigilancia_em_saude/controle_de_zoonoses/#

Dantas-Torres F, Solano-Gallego L, Baneth G, Ribeiro VM, de Paiva-Cavalcanti M, Otranto D. (2012) Canine leishmaniosis in the Old and New Worlds: unveiled similarities and differences. Trends in Parasitology, 28(12),531–538.

De Oliveira, LS, Rodrigues FM, de Oliveira FS, Mesquita PRR, Leal DC, Alcântara AC, Souza BM, Franke CR, Pereira PAP, de Andrade JB. (2008) Headspace solid phase microextraction/gas chromatography–mass spectrometry combined to chemometric analysis for volatile organic compounds determination in canine hair: A new tool to detect dog contamination by visceral leishmaniasis. Journal of Chromatography B. 875(2),392–398.

Drea CM, Boulet M, Delbarco-Trillo J, Greene LK, Sacha CR, Goodwin TE, Dubay GR. (2013) The “Secret” in Secretions: Methodological Considerations in Deciphering Primate Olfactory Communication. Am. J. Primatol. 75:621–642, https://doi.org/10.1002/ajp.22143

Ezenwa VO, Williams AE. (2014) Microbes and animal olfactory communication: Where do we go from here? BioEssays News Rev Mol Cell Dev Biol. 36(9):847–54. https://doi.org/10.1002/bies.201400016

Figueiredo FB, de Vasconcelos TCB, de Fátima Madeira M, Menezes RC, Maia-Elkhoury ANS, Marcelino AP, Werneck GL (2018) Validation of the Dual-path Platform chromatographic immunoassay (DPP^®^ CVL rapid test) for the serodiagnosis of canine visceral leishmaniasis. Mem Inst Oswaldo Cruz, Rio de Janeiro, Vol. 113(11): e180260.

Fraga DBM, Pacheco LV, Borja LS, Tuy PGdSE, Bastos LA, Solcà MdS, Amorim LDA, Veras PST (2016) The Rapid Test Based on Leishmania infantum Chimeric rK28 Protein Improves the Diagnosis of Canine Visceral Leishmaniasis by Reducing the Detection of False-Positive Dogs. PLoS Negl Trop Dis 10(1): e0004333. doi:10.1371/journal.pntd.0004333

Fraley, C. and A. E. Raftery (1999). “MCLUST: Software for model-based cluster analysis.” Journal of Classification 16(2): 297–306.

Fraley, C. and A. E. Raftery (2007). “Model-based methods of classification: Using the mclust software in chemometrics.” Journal of Statistical Software 18(6).

Greer CE, Peterson SL, Kiviat NB, Manos MM. (1991) PCR amplification from paraffin-embedded tissues. Effects of fixative and fixation time. Am. J. Clin. Pathol. 95, 117–124.

Grimaldi Jr G, Teva A, Ferreira AL, dos Santos CB, Pinto IdeS, de Azevedo CT, Falqueto A (2012) Evaluation of a novel chromatographic immunoassay based on Dual-Path Platform technology (DPP^®^ CVL rapid test) for the serodiagnosis of canine visceral leishmaniasis. Transactions of the Royal Society of Tropical Medicine and Hygiene 106;54–59

Hamilton, J.G.C. and Hurd, H. (2002) Parasite manipulation of vector behavior. In: The Behavioural Ecology of Parasites. (eds. Lewis, E.E., Campbell, J.F. and Sukhdeo, M.V.K.) 384pp CABI Publishing, Wallingford, pp. 259–281.

Kolk A, Hoelscher M, Maboko L, Jung J, Kuijper S, Cauchi M, Bessant C, Van Beers S, Dutta R, Gibson T, Reither K. (2010) Electronic-Nose Technology Using Sputum Samples in Diagnosis of Patients with Tuberculosis. Journal of Clinical Microbiology, 48(11),4235–4238 doi:10.1128/JCM.00569-10

Kücklich M, Möller M, Marcillo A, Einspanier A, Weiß BM, Birkemeyer C, Birkemeyer C, Widdig A. (2017) Different methods for volatile sampling in mammals. PLoS ONE 12(8): e0183440. https://doi.org/10.1371/journal.pone.0183440

Kumar S, Huang J, Abbassi-Ghadi N, Mackenzie HA, Veselkov KA, Hoare JM, Lovat LB, Španěl P, Smith D, Hanna GB. (2015). Mass Spectrometric Analysis of Exhaled Breath for the Identification of Volatile Organic Compound Biomarkers in Esophageal and Gastric Adenocarcinoma. Ann Surg, 262(6), 981–990.

Lacroix R, Mukabana WR, Gouagna LC, Koella JC (2005) Malaria Infection Increases Attractiveness of Humans to Mosquitoes. PLoS Biol 3(9): e298. https://doi.org/10.1371/journal.pbio.0030298

Liddell K. (1976) Smell as a diagnostic marker. Postgraduate Medical Journal 52,136–138.

Magalhães-Junior JT, Mesquita PRR, dos Santos Oliveira WF, Oliveira FS, Franke CR, Rodrigues FM, de Andrade JB, Barrouin-Melo SM (2014) Identification of biomarkers in the hair of dogs: new diagnostic possibilities in the study and control of visceral leishmaniasis. Anal Bioanal Chem. 406:6691–6700.

Malaquias LC, do Carmo Romualdo R, do Anjos JB, Jr., Giunchetti RC, Correa-Oliveira R, Reis AB. (2007) Serological screening confirms the re-emergence of canine leishmaniosis in urban and rural areas in Governador Valadares, Vale do Rio Doce, Minas Gerais, Brazil. Parasitol Res. 100(2):233–9. doi: 10.1007/s00436-006-0259-z. PubMed PMID: 16941189.

Mancianti F, Gramiccia M, Gradoni L, Pieri S. (1988) Studies on canine leishmaniasis control. 1. Evolution of infection of different clinical forms of canine leishmaniasis following antimonial treatment. Trans R Soc Trop Med Hyg. 1988;82(4)

Mary C, Faraut F, Lascombe L, Dumon H. (2004) Quantification of Leishmania infantum DNA by a real-time PCR assay with high sensitivity. J Clin Microbiol 42: 5249–5255.

MS/SVS/DVE (2014) Ministério da Saúde/Secretaria de Vigilância em Saúde/Departamento de Vigilância Epidemiológica. Manual de vigilância e controle da leishmaniose visceral. Série A. Normas e Manuais Técnicos. Brasília: MS; 120 pp.

Nakhleh, MK, Amal H, Jeries R, Broza YY, Aboud M, Gharra A, Ivgi H, Khatib S, Badarneh S, Har-Shai L, Glass-Marmor L, Lejbkowicz I, Miller A, Badarny S, Winer R, Finberg J, Cohen-Kaminsky S, Perros F, David Montani D, Barbara Girerd B, Garcia G, Simonneau G, Nakhoul F, Baram S, Salim R, Hakim M, Gruber M, Ronen O, Marshak T, Doweck I, Nativ O, Bahouth Z, Shi D, Zhang W, Hua Q, Pan Y, Tao L, Liu H, Karban A, Koifman E, Rainis T, Skapars R, Sivins A, Ancans G, Liepniece-Karele I, Kikuste I, Lasina I, Tolmanis I, Johnson D, Millstone S, Fulton J, Wells J, Wilf L, Humbert M, Leja M, Peled N, Haick H. (2017) Diagnosis and classification of 17 diseases from 1404 subjects via pattern analysis of exhaled molecules. ACS Nano, 11, 112−125 DOI: 10.1021/acsnano.6b04930

Nevatte TM, Ward RD, Sedda L. Hamilton JGC. (2017) After infection with Leishmania infantum, Golden Hamsters (Mesocricetus auratus) become more attractive to female sand flies (Lutzomyia longipalpis). Scientific Reports doi:10.1038/s41598-017-06313-w (http://rdcu.be/ulYY)

O’Hagan, A., T. B. Murphy and I. C. Gormley (2012). “Computational aspects of fitting mixture models via the expectation-maximization algorithm.” Computational Statistics & Data Analysis 56(12): 3843–3864.

OPAS/WHO (2017) Organización Panamericana de la Salud/World Health Organization. Leishmaniasis. Informe epidemiológico en las Américas. No. 5. Washington: OPAS; Available from: http://iris.paho.org/xmlui/bitstream/handle/123456789/34111/in-forme_leishmaniasis_5_spa.pdf?sequence=5&isAllowed=y.

O’Shea B, Rebollar-Tellez E, Ward RD, Hamilton JGC, EI Naiem D, Polwart A. (2002) Enhanced sandfly attraction to Leishmania-infected hosts. Trans. Royal Soc. Trop. Med. & Hyg. 96, 117–118.

Peeling RW, Holmes KK, Mabey D, Ronald A. (2006) Rapid tests for sexually transmitted infections (STIs): the way forward Sexually Transmitted Infections 82:v1–v6. doi: 10.1136/sti.2006.024265

Quinnell RJ, Courtenay O (2009) Transmission, reservoir hosts and control of zoonotic visceral leishmaniasis. Parasitology 136: 1915–1934.

R Core Team (2018). R: A language and environment for statistical computing Vienna, Austria, R Foundation for Statistical Computing.

Robinson A, Busula AO, Voets MA, Beshir KB, Caulfield JC, Powers SJ, Verhulst NO, Winskill P, Muwanguzi J, Birkett MA, Smallegange RC, Masiga DK, Mukabana WR, Sauerwein RW, Sutherland CJ, Bousema T, Pickett JA, Takken W, Logan JG, de Boer JG. (2018) Plasmodium-associated changes in human odor attract mosquitoes. PNAS 115(18), E4209–E4218 https://doi.org/10.1073/pnas.1721610115

Salomón OD, Feliciangeli MD, Quintana MG, Afonso MMS, Rangel EF. (2015) Lutzomyia longipalpis urbanisation and control. Mem Inst Oswaldo Cruz. 2015; 110(7): 831–46.

Scrucca, L., M. Fop, T. B. Murphy and A. E. Raftery (2016). “mclust 5: Clustering, Classification and Density Estimation Using Gaussian Finite Mixture Models.” R Journal 8(1): 289–317.

Spanakos G, Piperaki ET, Menounos PG, Tegos N, Flemetakis A, Vakalis NC (2008) Detection and species identification of Old World Leishmania in clinical samples using a PCR-based method. Transactions of the Royal Society of Tropical Medicine and Hygiene. 102, 46—53.

Smith D and Španěl P. (2016). Status of selected ion flow tube MS: accomplishments and challenges in breath analysis and other areas. Bioanalysis. 8(11), 1183–1201.

von Zuben APB, Donalísio MR. (2016) Dificuldades na execução das diretrizes do Programa de Vigilância e Controle da leishmaniose visceral em grandes municípios brasileiros. Cad Saude Publica. 32(6), 1–11.

Werneck GL. (2010) Geographic spread of visceral leishmaniasis in Brazil. Cad Saude Publica. 26(4), 645-. PubMed PMID: WOS:000278148700001.

