## Supplementary Material for "Volatile organic chemical analyser (eNose) diagnosis of dogs naturally infected with visceral leishmaniasis in Brazil"

**Supplementary Material 1.**

Rationale for the Data Analysis Approach.

Clustering

Rather than use conventional clustering algorithms (where the number of clusters is known) to analyse the data, we assumed that the data originated from an unknown number of clusters (Fraley and Raftery 1999) which (Gaussian) distributions are defined by several parameters (i.e. shape, orientation and volume). The MCLUST program, which is based on mixture modelling, has the advantage to provide an objective estimate of the number of clusters in the data.

The analysis was initiated by using an agglomerative hierarchical clustering method (i.e. a clustering method that groups records in a hierarchy or tree topped by a common ancestor) for multivariate data; while fitting was performed by the expectation-maximization (EM) algorithm (O'Hagan et al., 2012). The MCLUST program then searched for the optimal model (in terms of Bayesian Information Criterion (BIC) or Integrated Complete-data Likelihood criterion (ICL)) for the clustering of data among models with varying shape, orientation and volume of the distributions, as well as the number of components and substructures.
The mixture model that we used can be formally described by the following equation:

$$f(.) = w_1 f(.|\psi_1) + w_2 f(.|\psi_2) + \cdots + w_m f(.|\psi_m) \qquad [1]$$

where the probability density function, *f(.)*, is a linear combination of *m* probability density functions on the form $f(.|\psi)$, which for a Gaussian mixture model represent a Normal probability density function with parameter $\psi$:

$$f_k(.|\psi_k) \sim N(\mu_k, \Sigma_k) \qquad [2]$$

In the Gaussian Mixture Model, clusters are ellipsoidal with mean $\mu$, and volume, shape and orientation determined by the covariance matrix $\Sigma$. Finally, *m* is the number of mixture components (or classes) and *w* represents the relative frequency of class *k* (from 1 to *m*) in the data.

In MCLUST, each covariance matrix is decomposed by eigen decomposition in the form:

$$\Sigma_k = \lambda_k \mathrm{Q}_k \mathrm{D}_k Q_k^{-1} \qquad [3]$$

where for the k class, $Q$ is the orthogonal matrix of eigenvectors – which determines the orientation of the principal components, $D$ is a diagonal matrix whose elements are proportional to the eigenvalues of $\Sigma$ – i.e. the shape of the distribution, and $\lambda$ is a scalar – i.e. the volume of the distribution.  For multivariate data, 14 models with varying distribution (spherical, diagonal and ellipsoidal), volume (equal or variable across classes), shape (equal or variable across classes) and orientation (equal or variable across classes) are compared in MCLUST.

The EM algorithm computed the maximum likelihood estimators of the mixture model (with eigen-decomposed covariance). Convergence of the EM algorithm was checked when the change in log-likelihood or the relative change in log-likelihood was sufficiently small (O'Hagan, Murphy et al. 2012). Computation of the Bayesian Information Criterion (BIC) values was used to determine the form of the covariance structure and the number of components, which in turn determined the optimal number of clusters (O'Hagan, Murphy et al. 2012). Since BIC is the value of the maximized log likelihood penalised for the number of parameters in the model, it allowed the comparison of models with different parameterizations and/or different numbers of clusters (Andrews and McNicholas 2012). The closer the BIC value was to zero, the stronger the evidence for the model.

Finally, we tested the hypothesis of equality between two classes, using a bootstrapped likelihood ratio test: for each bootstrapped sample, the likelihoods with *m* and *m** classes were calculated followed by the likelihood ratio. The *P*-value was obtained by comparing the number of times that the bootstrapped likelihood ratio was larger than the observed (in the original data) likelihood ratio (Scrucca, Fop et al. 2016) over T replications.

Supervised Classification (discriminant analysis)

Once the optimal model and number of classes were decided (in terms of BIC); training errors, training sensitivity and training specificity were calculated. We used an Eigen Decomposition Discriminant Analysis (EDDA) within MCLUST which is based on equation [3] (Fraley and Raftery 2007). Therefore, when classes (and therefore their covariances and means) were obtained, the discriminant analysis was applied to find the separation plane(s) between classes. When this plane did not perfectly separate two classes training errors (misclassifications) resulted. In addition, the uncertainty, i.e. 1 minus the (sum of) probability that a member belonged a to different class(s), estimated the robustness of the classification since the classification probabilities were more informative than the assigned cluster labels.

One out-of-sample cross-validation (CV) and confounder cross validation (CCV).

To evaluate the predictivity of the clustering model we employed a traditional cross-validation method to predict out of sample membership (CV, results in the first two columns of Table 5). In addition, we have developed a CCV, based on a class-member permutation, to evaluate the capacity of the model to identify false members (or confounders) within a class. In practice, 10% of infected dog measurements were classified uninfected (false uninfected), while 10% of uninfected dogs measurements were classified infected (false infected). Then the full discriminatory analysis is repeated. We finally evaluated how many of the changed labels were correctly predicted (i.e. false uninfected predicted as real infected and false infected predicted as real uninfected) (results in the last two columns of Table 5).

Selecting important variables.

To compare the importance of each variable in determining the number of sub-classes (in both infected and uninfected dogs) a clustering with permuted variable values was performed. In other words, the tested (for importance) variable values are permuted, while the rest of the variables are kept unchanged, and clustering is performed. Then it is checked if the optimal number of sub-classes is larger or smaller than the one obtained with the un-permuted variable. The permutation is repeated 999 times, and a p-value produced as the number of times that the optimal number of sub-classes obtained by permutation is larger or shorter than the observed one, divided by 1000. Larger is the *P*-value, more important the variable is.

Reference not in the main text.

O'Hagan, A., T. B. Murphy and I. C. Gormley (2012). "Computational aspects of fitting mixture models via the expectation-maximization algorithm." Computational Statistics & Data Analysis **56**(12): 3843-3864.

Supplementary Table S1. Outcome of the mixture model analysis showing the top three models for the 2017 uninfected and infected dog data.

| Classes | Model 1 | Model 2 | Model 3 |
|---|---|---|---|
| Uninfected | EEE 1 class (-90584) | EEV 1 class (-90584) | EVE 1 class (-90584) |
| Infected | VVI 3 classes (-25125) | VEI 4 classes (-25169) | VEI 3 classes (-25180) |

Bayesian Information Criterion (BIC) values are shown within brackets. The closer the BIC value is to zero, the stronger the evidence for the model.

Supplementary Table S2. Outcome of the mixture model analysis showing the top three models for the 2018 uninfected and infected dog data.

| Classes | Model 1 | Model 2 | Model 3 |
|---|---|---|---|
| Uninfected | EEE 2 classes (-93455) | EEE 3 classes (-93621) | EEE 4 classes (-93884) |
| Infected | EEE 6 classes (-212458) | EEV 6 classes (-212628) | EVE 6 classes (-212988) |

Bayesian Information Criterion (BIC) values are shown within brackets. The closer the BIC value is to zero, the stronger the evidence for the model.

Supplementary Table S3. Relative Importance of different sensor variables in the contribution to the clustering observed in 2017 data.

| variable | *P*-value | variable | *P*-value | variable | *P*-value | variable | *P*-value |
|---|---|---|---|---|---|---|---|
| F3.Abs.5 | 0.93 | F2.Area.12 | 0.03 | F3.Abs.23 | 0.01 | F2.Area.8 | 0 |
| F1.Div.20 | 0.92 | F1.Div.12 | 0.03 | F1.Div.10 | 0.01 | F3.Abs.11 | 0 |
| F4.Des.8 | 0.07 | F2.Area.19 | 0.03 | F4.Des.4 | 0.01 | F3.Abs.22 | 0 |
| F1.Div.5 | 0.06 | F2.Area.15 | 0.03 | F2.Area.22 | 0.01 | F3.Abs.4 | 0 |
| F1.Div.19 | 0.06 | F4.Des.23 | 0.02 | F4.Des.22 | 0.01 | F2.Area.24 | 0 |
| F1.Div.13 | 0.05 | F1.Div.18 | 0.02 | F2.Area.17 | 0.01 | F4.Des.5 | 0 |
| F1.Div.16 | 0.05 | F3.Abs.24 | 0.02 | F4.Des.12 | 0.01 | F4.Des.1 | 0 |
| F1.Div.11 | 0.05 | F1.Div.7 | 0.02 | F4.Des.19 | 0.01 | F4.Des.6 | 0 |
| F4.Des.24 | 0.05 | F4.Des.16 | 0.02 | F3.Abs.17 | 0.01 | F4.Des.21 | 0 |
| F1.Div.15 | 0.05 | F3.Abs.13 | 0.02 | F4.Des.9 | 0.01 | F2.Area.16 | 0 |
| F1.Div.21 | 0.04 | F3.Abs.1 | 0.02 | F4.Des.17 | 0.01 | F3.Abs.19 | 0 |
| F2.Area.10 | 0.04 | F4.Des.10 | 0.02 | F1.Div.3 | 0.01 | F4.Des.13 | 0 |
| F2.Area.5 | 0.04 | F2.Area.20 | 0.02 | F2.Area.1 | 0.01 | F2.Area.23 | 0 |
| F3.Abs.9 | 0.03 | F1.Div.6 | 0.02 | F2.Area.3 | 0.01 | F2.Area.21 | 0 |
| F3.Abs.7 | 0.03 | F1.Div.17 | 0.02 | F4.Des.3 | 0.01 | F2.Area.7 | 0 |
| F1.Div.8 | 0.03 | F4.Des.20 | 0.02 | F1.Div.9 | 0 | F4.Des.7 | 0 |
| F4.Des.11 | 0.03 | F2.Area.6 | 0.02 | F3.Abs.21 | 0 | F1.Div.1 | 0 |
| F1.Div.24 | 0.03 | F2.Area.9 | 0.02 | F3.Abs.6 | 0 | F2.Area.4 | 0 |
| F3.Abs.18 | 0.03 | F3.Abs.15 | 0.02 | F3.Abs.12 | 0 | F2.Area.18 | 0 |
| F1.Div.4 | 0.03 | F3.Abs.10 | 0.01 | F3.Abs.16 | 0 | F2.Area.13 | 0 |
| F2.Area.11 | 0.03 | F1.Div.23 | 0.01 | F3.Abs.8 | 0 | F3.Abs.3 | 0 |
| F1.Div.22 | 0.03 | F3.Abs.20 | 0.01 | F4.Des.18 | 0 | F4.Des.15 | 0 |

Abs = absorbance, Des = desorbance, Area = area under the curve, Div = divergence.

Supplementary Table S4. Relative Importance of different sensor variables in the contribution to the clustering observed in 2018 data.

| variable | *P*-value | variable | *P*-value | variable | *P*-value | variable | *P*-value |
|---|---|---|---|---|---|---|---|
| F3.Abs.17 | 0.99 | F3.Abs.21 | 0.62 | F4.Des.13 | 0.52 | F3.Abs.5 | 0.42 |
| F4.Des.21 | 0.98 | F3.Abs.8 | 0.61 | F4.Des.20 | 0.52 | F4.Des.23 | 0.42 |
| F4.Des.9 | 0.95 | F4.Des.14 | 0.61 | F1.Div.3 | 0.51 | F2.Area.5 | 0.41 |
| F4.Des.4 | 0.94 | F1.Div.21 | 0.6 | F1.Div.22 | 0.51 | F4.Des.22 | 0.41 |
| F4.Des.6 | 0.91 | F4.Des.18 | 0.6 | F2.Area.20 | 0.51 | F3.Abs.7 | 0.4 |
| F2.Area.6 | 0.82 | F3.Abs.10 | 0.58 | F3.Abs.15 | 0.51 | F2.Area.2 | 0.39 |
| F4.Des.19 | 0.81 | F1.Div.4 | 0.57 | F2.Area.15 | 0.5 | F2.Area.17 | 0.35 |
| F3.Abs.1 | 0.79 | F2.Area.1 | 0.57 | F1.Div.5 | 0.49 | F3.Abs.3 | 0.35 |
| F4.Des.8 | 0.78 | F1.Div.10 | 0.56 | F2.Area.4 | 0.49 | F1.Div.20 | 0.33 |
| F2.Area.9 | 0.77 | F2.Area.3 | 0.56 | F2.Area.14 | 0.49 | F3.Abs.4 | 0.33 |
| F2.Area.21 | 0.77 | F2.Area.23 | 0.56 | F4.Des.15 | 0.47 | F1.Div.18 | 0.3 |
| F1.Div.19 | 0.74 | F3.Abs.6 | 0.56 | F4.Des.16 | 0.47 | F4.Des.3 | 0.3 |
| F3.Abs.13 | 0.74 | F1.Div.11 | 0.55 | F1.Div.16 | 0.46 | F4.Des.17 | 0.29 |
| F2.Area.22 | 0.69 | F2.Area.13 | 0.55 | F2.Area.16 | 0.46 | F1.Div.6 | 0.27 |
| F1.Div.17 | 0.68 | F3.Abs.14 | 0.55 | F2.Area.18 | 0.46 | F1.Div.15 | 0.27 |
| F2.Area.10 | 0.68 | F4.Des.2 | 0.55 | F1.Div.13 | 0.45 | F4.Des.1 | 0.21 |
| F4.Des.7 | 0.67 | F2.Area.8 | 0.54 | F1.Div.14 | 0.45 | F3.Abs.2 | 0.17 |
| F1.Div.9 | 0.66 | F3.Abs.22 | 0.54 | F2.Area.11 | 0.45 | F3.Abs.16 | 0.13 |
| F3.Abs.20 | 0.66 | F1.Div.7 | 0.53 | F3.Abs.9 | 0.44 | F1.Div.1 | 0.1 |
| F3.Abs.19 | 0.65 | F4.Des.10 | 0.53 | F4.Des.11 | 0.44 | F4.Des.5 | 0.05 |
| F3.Abs.11 | 0.64 | F1.Div.23 | 0.52 | F1.Div.2 | 0.43 | F3.Abs.18 | 0.04 |
| F3.Abs.23 | 0.64 | F2.Area.19 | 0.52 | F2.Area.7 | 0.43 | F1.Div.8 | 0.02 |

Abs = absorbance, Des = desorbance, Area = area under the curve, Div = divergence.
